# Circulating Microplastics as Acute Triggers of Platelet Activation and Coagulation: Implications for Cardiovascular Risk

**DOI:** 10.64898/2025.12.05.692697

**Authors:** A. Trostchansky, M. Alarcón

**Affiliations:** Departamento de Bioquímica and Center for Free Radical and BiomedicalResearch (CEINBIO), Facultad de Medicina, Universidad de la República, Montevideo 11800, Uruguay; Thrombosis Research Center and Healthy Aging, Universidad de Talca, 2 Norte 685, Talca, Chile; Department of Clinical Biochemistry and Immunohematology, Faculty of Health Sciences, Universidad de Talca, 2 Norte 685, Talca, Chile

**Author notes:** Corresponding authors: Andrés Trostcansky, PhD, Facultad de Medicina, Universidad de la República, P.O. Box 11800, Montevideo, Uruguay, Marcelo Alarcón, PhD, Faculty of Health Sciences, Universidad de Talca, Talca, Chile, P.O. Box 747, Talca, Chile.

## Abstract

**Background:** Microplastics and nanoplastics (MPs/NPs) have recently been detected in human blood and vascular tissues, yet their direct effects on thrombosis remain poorly defined. Given the central role of platelets in atherothrombotic disease, understanding how circulating NPs exposure influences platelet function is critical for cardiovascular health.

**Methods:** Washed platelets and citrated whole blood from healthy volunteers were exposed to fluorescent carboxylated polystyrene nanoplastics (PS-NPs; 100 nm). PS-NPs association, internalization, and activation were quantified by flow cytometry (side scatter, forward scatter, PS-NPs fluorescence, and CD63). Fluorescence microscopy visualized the PS-NPs uptake kinetics. A whole-blood coagulation assay assessed PS-NPs-induced clot formation under varying Ca²⁺ concentrations.

**Results:** PS-NPs rapidly associated with human platelets in a concentration and time-dependent manner, with near-maximal internalization achieved within 10 minutes. PS-NPs uptake induced marked structural remodeling (increased FSC/SSC), pseudopod formation, and concentration-dependent CD63 externalization, indicative of robust platelet activation comparable to that induced by thrombin stimulation. PS-NPs association remained non-saturable across tested doses (0.3–20 μg), suggesting high-capacity, non-specific uptake mechanisms. In whole blood, PS-NPs induced dense fibrin clot formation exclusively under Ca²-permissive conditions, and PS-NPs were incorporated into fibrin networks, consistent with their role as catalytic pro-coagulant surfaces. Supernatant fluorescence confirmed PS-NPs sequestration into clots.

**Conclusions:** PS-NPs rapidly bind and activate human platelets, promote pro-coagulant platelet phenotypes, and integrate into fibrin-rich thrombi. These findings provide mechanistic evidence that circulating MPs may function as previously unrecognized environmental cardiovascular risk factors capable of acutely enhancing thrombotic potential. Defining exposure thresholds and *in vivo* relevance is urgently needed to assess the cardiovascular impact of the rising human NPs burden.

**Clinical Perspective:** *What Is New?:*

- Circulating nanoplastics rapidly bind to human platelets through high-capacity, non-specific membrane interactions, triggering cytoskeletal remodeling, granule release, and activation pathways similar to classical agonists such as thrombin.
- Nanoplastics serve as catalytic pro-coagulant surfaces that accelerate fibrin polymerization and become structurally embedded within thrombi, providing a direct mechanistic link between environmental microplastic exposure and thrombogenesis.
- Rapid, non-saturable platelet uptake suggests that rising global microplastic loads may proportionally increase human thrombotic susceptibility.

*What Are the Clinical Implications?:*

- Nanoplastic-induced platelet activation and clot formation identify microplastics as previously unrecognized, modifiable environmental cardiovascular risk factors with mechanistic plausibility for contributing to myocardial infarction, stroke, and microvascular thrombosis.
- Populations with heightened platelet reactivity, endothelial dysfunction, or chronic inflammatory states may be particularly vulnerable to thrombotic events triggered by acute or cumulative nanoplastic exposure.
- Public health efforts aimed at reducing environmental microplastic contamination and establishing exposure limits may become necessary components of cardiovascular disease prevention strategies.

## Introduction

Since the mid-twentieth century, global plastic production has increased exponentially, resulting not only in large-scale plastic waste but also in the generation of microscopic plastic fragments known as microplastics (MPs), which are defined as particles smaller than 5 mm, while nanoplastics (NPs) are even smaller, defined as being less than 1 µm in size. Plastic fragments are now ubiquitous across marine, terrestrial, and atmospheric environments. They can originate as primary MPs, such as microbeads used in cosmetics and industrial precursors, or as secondary MPs, which are formed through degradation processes such as ultraviolet radiation, mechanical abrasion, and incomplete biodegradation of larger plastic debris^1–3^.

Human exposure to plastic fragments occurs through several routes, including ingestion of contaminated food and water, inhalation of airborne particles and fibers, and dermal contact. Biomonitoring studies have confirmed the presence of plastic fragments in human blood, urine, feces, placenta, and breast milk^1, 2^. The pervasive nature of plastic fragments, coupled with their ability to adsorb and release chemical additives such as acrylamide (AA) and bisphenol A (BPA), raises significant toxicological concerns^4, 5^.

The toxic effects of plastic fragments are influenced by both their physical characteristics and chemical composition. The small size and irregular morphology of plastic fragments facilitate tissue penetration and accumulation, potentially leading to inflammation and cytotoxicity. Additionally, the chemical additives associated with plastic fragments can induce oxidative stress, mitochondrial dysfunction, and disruption of the endocrine and immune systems^5–7^.AA and BPA are among the most commonly used plastic additives and have been detected in human tissues and fluids, where they are associated with neurotoxic, hepatotoxic, immunotoxic, and reproductive toxicities, primarily through the generation of reactive oxygen species (ROS) and oxidative stress^8^.

Furthermore, a recent study demonstrated that circulating plastic fragments are phagocytosed by immune cells in the bloodstream, leading to physical obstruction of cerebral capillaries, thrombus formation, and neurobehavioral abnormalities in murine models^9^. This finding underscores the capacity of plastic fragments to disrupt vascular physiology through both mechanical and cellular mechanisms. However, the direct pathways by which circulating plastic fragments interact with cellular blood components, such as platelets, to initiate acute prothrombotic events remain poorly defined.

Particularly concerning is the role of AA and BPA in modulating platelet function. Recent studies have shown that these compounds can enhance platelet activation and aggregation through oxidative stress pathways, contributing to a prothrombotic state that increases the risk of cardiovascular diseases (CVDs), such as myocardial infarction and stroke^5–7^. Platelets are key players in hemostasis, inflammation, and vascular pathology, positioning MPs and their associated chemical additives as emerging contributors to CVD risk.

Understanding the intricate pathways of human exposure, bioavailability, and the cellular mechanisms associated with plastic fragments and their chemical additives is essential for evaluating their effects on cardiovascular and overall systemic health. Emerging evidence highlights the pressing need for further investigation and regulatory measures to mitigate the health risks posed by ongoing environmental and human exposure to plastic fragments, AA, BPA, and similar compounds.

## Materials and Methods

Polystyrene nanoparticles (fluorescent 505/515 nm, carboxyl-modified; Thermo Fisher Scientific, USA) were used for all experiments. Sterile phosphate-buffered saline (PBS, 1×) was used for nanoparticle dilution, and particle dispersion was enhanced using a Clifton Ultrasonic Bath (Clifton, NJ, USA). Tyrode’s buffer without calcium, acid-citrate-dextrose (ACD), and 3.2% sodium citrate were used for platelet and whole blood handling. Human platelet identification and activation markers included anti-human CD41-APC-Cy7-A and CD63-PE antibodies (BD Biosciences, San Jose, CA, USA). Flow cytometric analyses were performed using a BD FACS Lyric cytometer, and microscopy was conducted using an Axio Examiner Z1 fluorescence microscope, equipped with an Axiocam 506 camera. Image processing and quantitative analyses were performed using ImageJ software (NIH; version 1.26t). All centrifugations were conducted using an Eppendorf 5804 centrifuge (Hamburg, Germany). Platelet counts were obtained using a Mindray BC-3000 Plus hematology analyzer. Calcium chloride (CaCl₂, 100 mM) was used to induce coagulation in whole-blood assays.

### Nanoparticle Preparation

Polystyrene nanoparticles (PS-NPs) of 100 nm diameter (fluorescent beads, 505/515 nm), with an anionic surface chemistry (carboxyl-modified, catalog: F8803, Thermo Fisher Scientific, USA), were used in this study as representative particles of plastic fragment contaminants. PS-NPs were diluted in sterile-filtered phosphate-buffered saline (PBS, 1X). To reduce particle agglomeration, the PS-NPs were sonicated for 1 minute using a Clifton Ultrasonic Bath (Clifton, NJ), followed by vortexing for 1 minute before dilution and just before use^10^.

### Platelet Isolation and Preparation

Platelet samples were prepared according to a standardized protocol established by our laboratory^5^, with minor modifications. Venous blood (10 mL) was collected from healthy volunteers into syringes containing acid-citrate-dextrose (ACD; 4:1 v/v). The blood was first centrifuged at 250 × g for 12 minutes (without brake or accelerator; Eppendorf 5804, Hamburg, Germany) to isolate the platelet-rich plasma (PRP) fraction. The PRP was then centrifuged at 900 × g for 8 minutes. The plasma supernatant was removed, and the platelet pellet was resuspended in Tyrode’s buffer without calcium, supplemented with ACD at a 5:1 v/v ratio. Platelet count was determined using a hematology counter (Mindray BC-3000 Plus, Japan). All donors were medication-free for at least 10 days before blood collection and exhibited no signs of bleeding disorders or acute illness. Samples exhibiting hemolysis or lipemia were excluded from analysis. Written informed consent was obtained from all participants before enrollment. The study protocol was approved by the Ethics Committee of the Universidad de Talca and complied with the principles of the Declaration of Helsinki (18th World Medical Assembly, Helsinki, 1964).

### Flow cytometric analysis

Flow cytometric analysis was performed using a BD FACS Lyric flow cytometer (BD Biosciences). Data collection was conducted until 10,000 total events were recorded. Forward scatter (FSC), side scatter (SSC), and fluorescence data were obtained using a logarithmic scale configuration. All buffers were double-filtered through a 0.22 μm filter. Representative data are presented in all figures. The platelet population (>99%) in washed platelets was identified using anti-human CD41-APC-Cy7-A antibody (BD Biosciences, San Jose, CA, USA). Data were expressed as mean fluorescence intensity (MFI)^11^. Washed platelets were treated with varying concentrations of PS-NPs (0–200 μg) or vehicle control for 10 minutes. Platelet activation was quantified using an antibody against CD63-PE, a marker of platelet degranulation. The potential interference of PS-NPs with the assay was assessed by measuring light scattering in the absence of cells (Supplementary Figure 1).

### Fluorescence Microscopy

For bead binding and localization studies, fluorescence microscopy images were captured using the 40x objective lens of an Axio Examiner Z1 fluorescence microscope (Zeiss). The images were subsequently analyzed using ImageJ software (version 1.26t, NIH). Image acquisition was carried out using an Axiocam 506 digital camera (Jena, Germany), and images were processed with ZEN2.3 Pro software (Jena, Germany).

### Whole Blood Coagulation Assay

For the whole blood coagulation assays, blood was collected from healthy human volunteers using 3.2% sodium citrate as an anticoagulant to preserve cellular and plasma components. Briefly, citrated whole blood was aliquoted into 500 μL samples, treated with 10 μg of PS-NPs or vehicle, and coagulation was induced by adding calcium chloride (100 mM CaCl₂) to achieve final concentrations of 2 mM, 4 mM, and 12 mM, with the 12 mM sample serving as the positive control. All samples were incubated at 37°C for 30 minutes. Clotting was visually assessed by gently inverting the tubes. Clots formed under the 4 mM Ca²⁺ + PS-NPs condition were subsequently analyzed for physical properties: clots were carefully isolated, blotted dry, and weighed using a precision balance. Arbitrary clot size (AU) was measured using ImageJ software.

### Statistical analysis

Platelet assays were performed using blood samples from six independent healthy volunteers, a sample size deemed appropriate based on the high reproducibility and low inter-individual variability of ex vivo platelet assays^5^. Each volunteer served as their own control, with platelet responses measured before and after exposure to PS-NPs, allowing for paired analysis to minimize variability and enhance statistical power. Data were analyzed using GraphPad Software version 6.0 (La Jolla, CA, USA). Two or more measurements were performed for each test, and data were expressed as mean ± standard error of the mean (SEM). Differences between groups were analyzed using Student’s t-test or one-way ANOVA, with p values < 0.05 considered statistically significant.

## Results and Discussion

### Nanoplastic Uptake by Washed Platelets is Time-Dependent and Kinetically Saturable

Platelets isolated from human donors were identified using flow cytometry with a sequential gating strategy (Supplementary Figure 1). Exposure to 2 μg of polystyrene nanoparticles (PS-NPs) led to a time-dependent increase in both fluorescence intensity and side scatter (SSC), indicating progressive internalization of the nanoparticles over 60 minutes (Figure 1A). The proportion of PS-NPs-positive events (R4) steadily increased, reaching a plateau after 10 minutes (Figure 1B). At higher PS-NP doses (200 μg), fluorescence saturation occurred within 10 minutes (Supplementary Figure 2), suggesting that platelet binding capacity reached its limit at supraphysiological concentrations, thus restricting further kinetic resolution.

**Figure 1.**
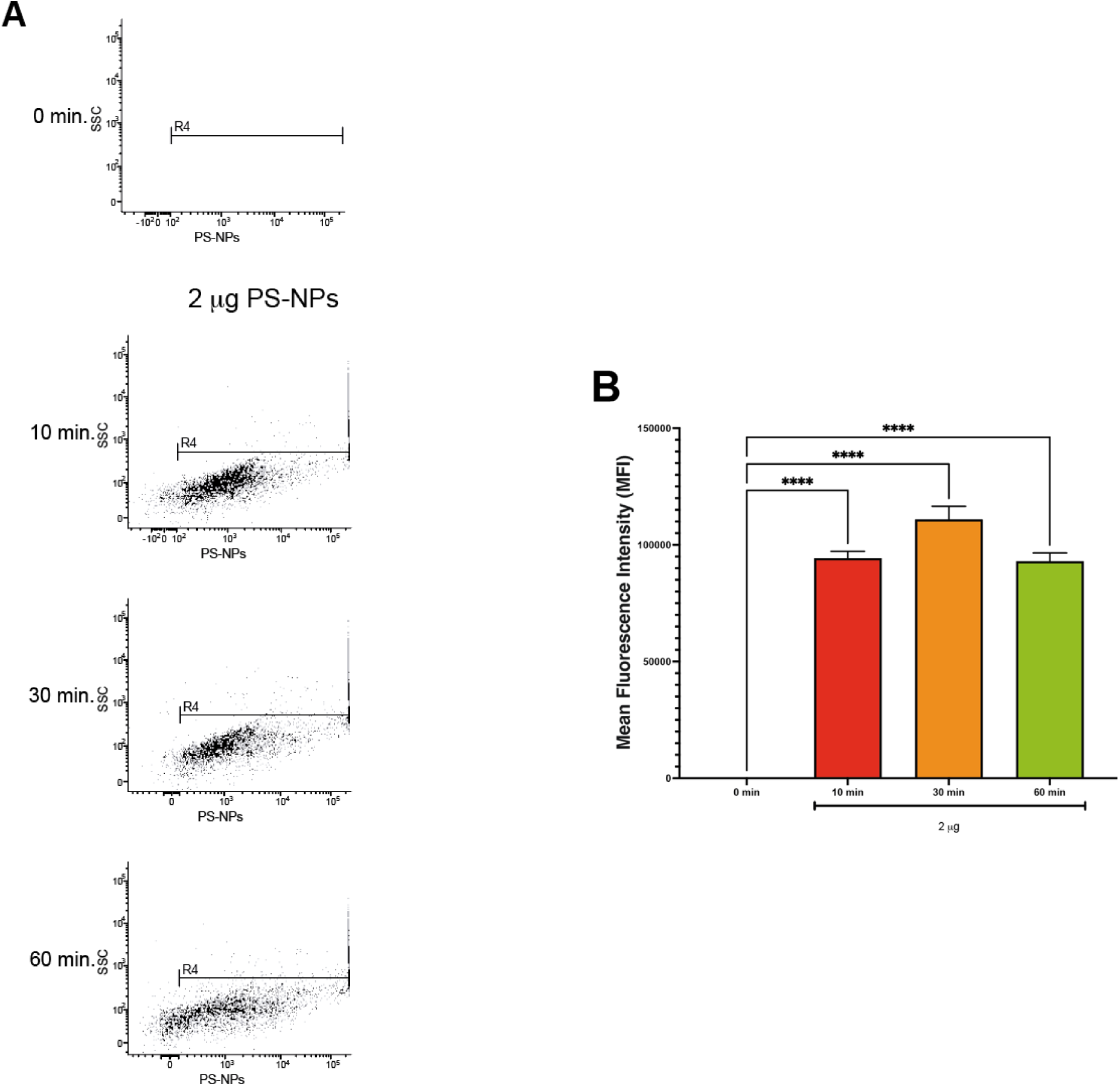
Kinetics of Polystyrene Nanoplastic (PS-NPs) Internalization by Washed Human Platelets. (A) Flow cytometry analysis of washed human platelets exposed to 2 μg of fluorescently labeled PS-NPs over time (0, 10, 30, and 60 minutes). Dot plots show the relationship between Side Scatter (SSC, Y-axis) and PS-NPs fluorescence intensity (X-axis, logarithmic scale). The 0-minute time point serves as an unexposed control, defining baseline fluorescence. Exposure to PS-NPs leads to a time-dependent increase in both fluorescence (PS-NPs uptake) and SSC (reflecting enhanced internal complexity), indicating PS-NPs internalization. Region R4 identifies platelets positive for PS-NPs fluorescence. (B) Mean Fluorescence Intensity (MFI) of washed platelets following PS-NPs exposure at 0, 10, 30, and 60 minutes. Data are presented as mean ± SEM; ****p < 0.0001.

The rapid kinetics of PS-NPs uptake can be attributed to their interaction with the highly dynamic, receptor-rich platelet plasma membrane, which facilitates quick binding and internalization^12^. This initial uptake likely involves physicochemical interactions, such as hydrophobic interactions and the formation of a protein corona^13^. The near-maximal uptake observed within 10 minutes supports the hypothesis that platelets may serve as rapid carriers for circulating PS-NPs, a finding with significant implications for thrombosis and hemostasis, given that PS-NPs have been detected in human blood and thrombotic lesions^1^. Such interactions may disrupt normal platelet function, potentially contributing to thrombotic risk^14^.

Quantification of mean fluorescence intensity (MFI) following 10, 30, or 60 minutes of exposure revealed a significant increase in MFI compared to baseline values (Figure 1B, p < 0.0001). However, no significant differences were observed between the 10, 30, and 60-minute time points, indicating that PS-NP uptake reached saturation by 10 minutes. Given this kinetic profile, a 10-minute exposure was selected for subsequent experiments to ensure maximal PS-NPs internalization while maintaining experimental efficiency.

These results confirm that platelets rapidly associate with PS-NPs, with near-maximal uptake achieved by 10 minutes. The concurrent rise in MFI and SSC supports the hypothesis that PS-NPs internalization is driven by high-affinity physicochemical interactions at the platelet membrane^12^. This rapid uptake may reflect a “pseudo-phagocytic” behavior of platelets^13^, reinforcing their potential role as carriers of environmental pollutants and implicating them in cardiovascular pathology.

### Nanoplastics Induce Platelet Activation and Morphological Changes

Exposure to 2 μg and 200 μgPS-NPs concentrations induced significant platelet activation, as evidenced by shifts in FSC and SSC (Figure 2), characteristic of platelet activation markers such as pseudopod formation, cytoskeletal remodeling, and granule mobilization. At the lower PS-NPs concentration (2 μg), approximately 70–76% of platelets shifted to the activated population (Figure 2A). At higher PS-NPs concentrations (200 μg, Figure 2B), fewer single platelets were activated (∼50%), but a broader distribution in FSC/SSC indicated more extensive morphological changes, including pseudopod extension and microaggregate formation^15, 16^.

**Figure 2.**
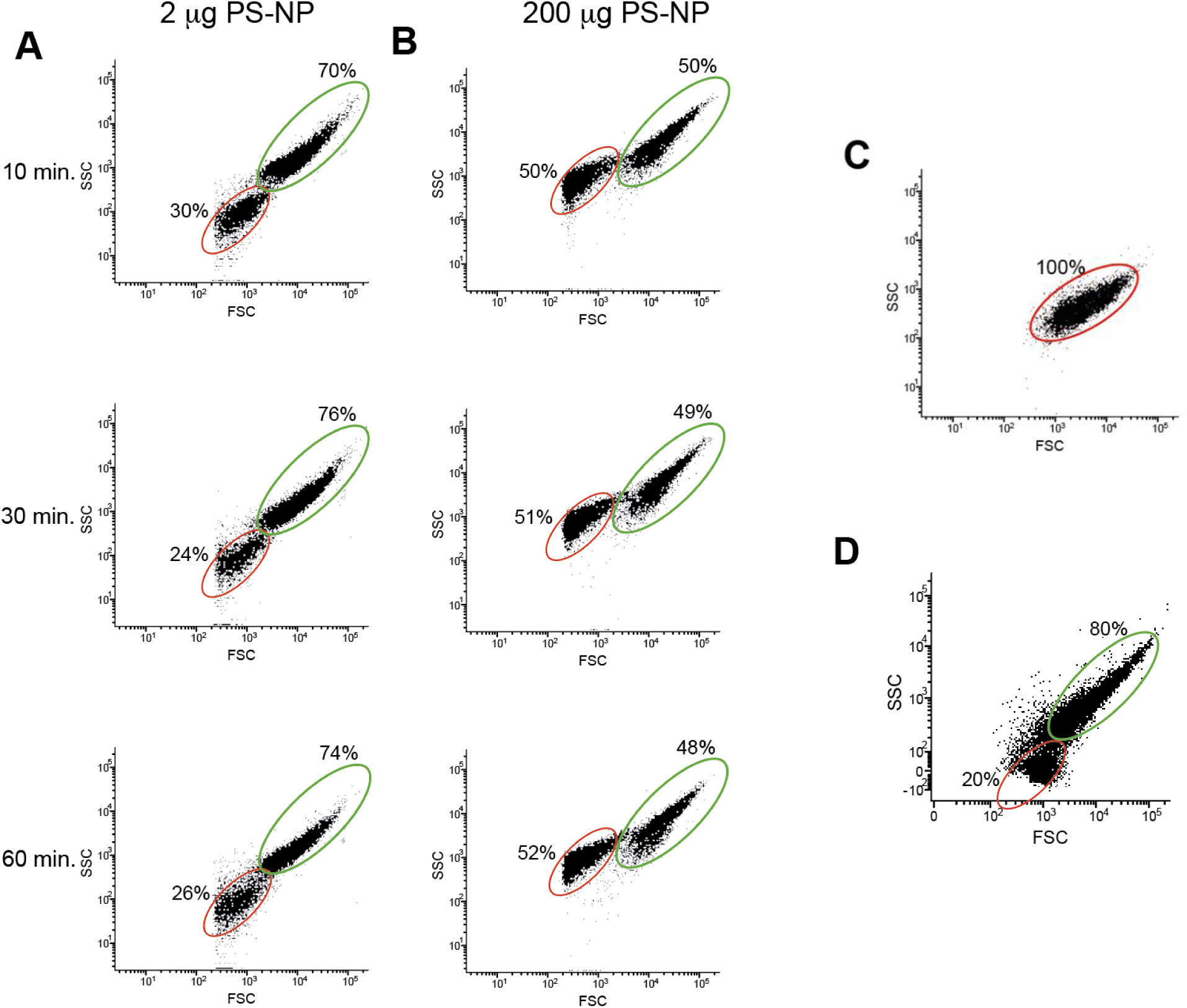
Effect of Polystyrene Nanoplastic (PS-NPs) Concentration and Thrombin on Platelet Morphology. (A-B) Flow cytometry analysis of washed human platelets exposed to varying concentrations of PS-NPs (2 μg in panel A, 200 μg in panel B) over a time course (10, 30, and 60 minutes). Dot plots show Forward Scatter (FSC; X-axis, reflecting cell size) versus Side Scatter (SSC; Y-axis, reflecting internal complexity). Two platelet populations are highlighted: resting (red gate) and activated-like (green gate), with the percentage of platelets in each population indicated. (C) Baseline scatter profile of unstimulated washed human platelets (0 minutes), showing the resting platelet population. (D) Positive control: Platelets stimulated with thrombin (10 U/mL) for 10 minutes. The thrombin-stimulated profile is used to delineate the activated-like population (green gate).

The PS-NPs-induced activation closely mirrored that of thrombin (Figure 2D), a physiological platelet activator, suggesting that PS-NPs may act through similar pathways, possibly involving receptors such as FcγRIIa or Toll-like receptors^17^. Furthermore, CD63 externalization confirmed that PS-NPs induced concentration-dependent degranulation, even at low PS-NPs concentrations (0.3 μg; Supplementary Figure 3). These data provide functional evidence that PS-NPs induce bona fide platelet activation, with implications for platelet aggregation and thrombus formation ^18^.

### Dose-Dependent Nanoplastic Association and Platelet Activation

To investigate the relationship between PS-NPs concentration and platelet responses, platelets were exposed to a range of PS-NPs doses (0.3–20 μg). MFI increased proportionally with PS-NPs concentration and did not reach saturation at any tested dose (Figure 3). The absence of a saturation plateau suggests that platelet-PS-NPs interactions occur through non-specific, high-capacity mechanisms, such as hydrophobic interactions and pseudo-phagocytosis^10, 19^. The progressive increases in SSC and fluorescence intensity observed with escalating PS-NPs doses (Supplementary Figures 4 and 5) indicate enhanced cytoskeletal reorganization and early pro-aggregatory behavior, both of which are critical steps in thrombosis^20^.

**Figure 3.**
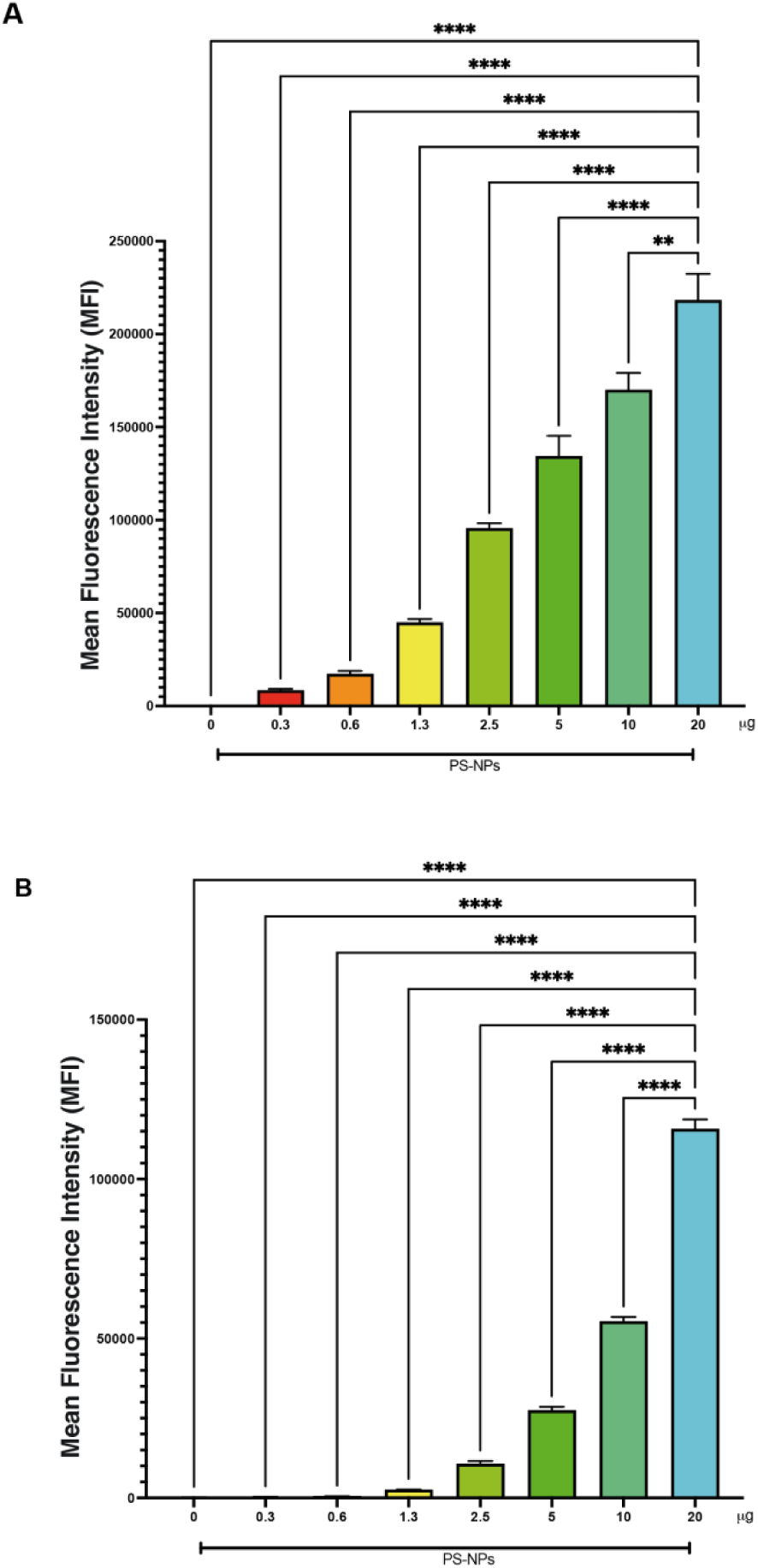
Dose-Dependent Polystyrene Nanoplastic (PS-NPs) Uptake in Washed Human Platelets. (A) Mean Fluorescence Intensity (MFI) of washed platelets exposed to varying concentrations of PS-NPs (0 to 20 μg) for a fixed 10-minute period (optimized in Figure 1). (B) MFI of washed platelets exposed to 0 to 20 μg of PS-NPs for 10 minutes. MFI quantifies the average cellular load of associated PS-NPs. Data are presented as mean ± SEM. A clear dose-dependent relationship is observed. ****: p < 0.0001; **: p < 0.01. Specific comparisons are indicated by brackets.

These findings suggest that PS-NPs accumulation in platelets scales with environmental exposure, providing a mechanistic basis for how increased PS-NPs burden may enhance platelet activation and elevate thrombotic risk. The strong, dose-dependent relationship between PS-NPs association and platelet activation highlights PS-NPs exposure as a potential driver of thrombotic events^21^. Our data demonstrate that PS-NPs act as potent activators of human platelets and catalysts of coagulation in vitro. These mechanistic observations align with recent *in vivo* findings showing that circulating PS-NPs particles are captured by neutrophils and macrophages, forming cellular aggregates that obstruct cerebral capillaries, reduce blood flow, and promote thrombosis^9^. It is plausible that the PS-NPs-induced platelet activation observed in our study may act synergistically with the cellular obstruction^9^, whereby activated platelets adhere to PS-NPs-laden immune cells lodged within the microvasculature, thereby amplifying thrombus formation.

### Nanoplastics Induce Coagulation in Whole Blood

The pro-thrombotic potential of PS-NPs was assessed in whole blood, where incubation with 10 μgPS-NPs for 30 minutes led to the formation of a dense fibrin clot. Microscopy confirmed the incorporation of PS-NPs into the fibrin network (Figure 4A), while quantification of the residual supernatant showed a significant decrease in MFI (Figure 4B), indicating efficient sequestration of PS-NPs into the clot. Coagulation was strictly dependent on Ca²⁺, with clot formation occurring only at 4 mM Ca²⁺ (Figure 4C). This suggests that PS-NPs do not directly induce coagulation but instead function as catalytic surfaces for the Ca²+-dependent assembly of coagulation factor complexes^22, 23^.

**Figure 4.**
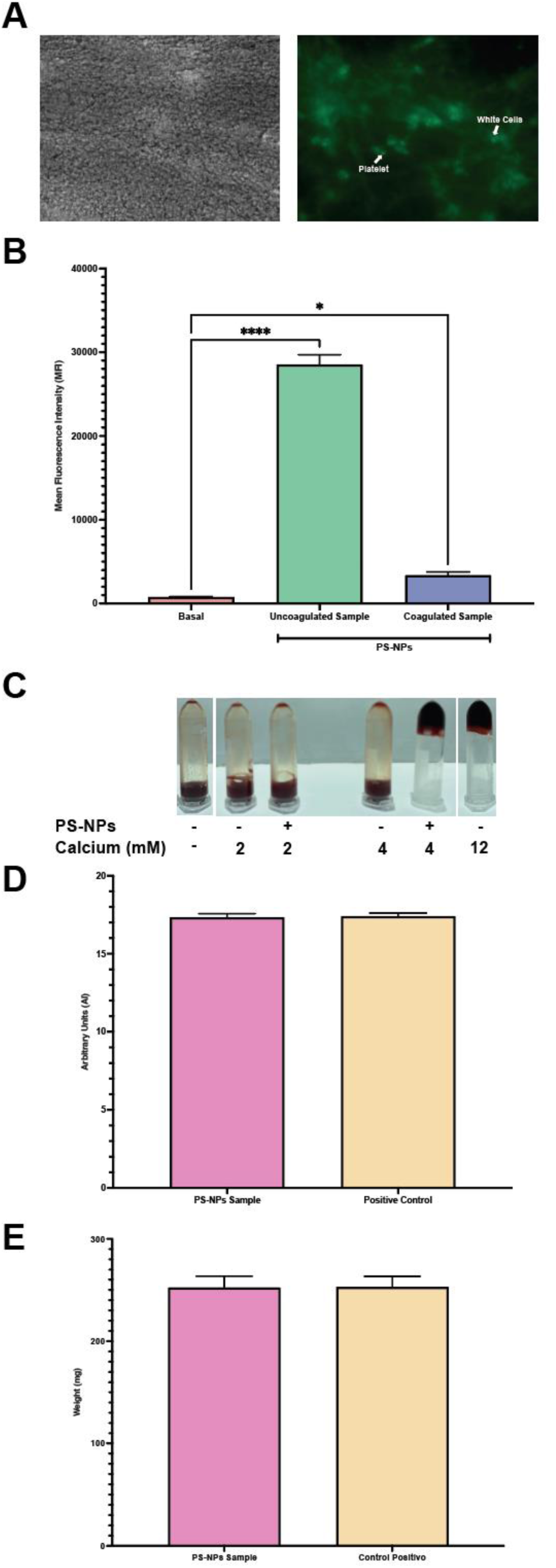
Polystyrene Nanoplastic (PS-NPs) Enhance Coagulation and Modulate Fibrin Clot Formation. (A) Microscopic analysis of whole blood clots after 30 minutes of incubation with 10 μg of PS-NPs. Left: Bright-field image shows dense fibrin mesh trapping blood cells. Right: Fluorescence image confirms PS-NPs incorporation within the clot matrix. (B) MFI of PS-NPs in the liquid phase after 30 minutes of whole blood incubation. The MFI of the Basal (no PS-NPs) sample is compared to the supernatant of uncoagulated and coagulated blood samples. The reduction in MFI in the coagulated sample indicates PS-NPs sequestration within the fibrin clot. (C) Test tube assay demonstrating calcium (Ca²⁺)-dependent pro-coagulant activity of PS-NPs. Whole blood samples were incubated with or without 10 μgPS-NPs at various Ca²⁺ concentrations (0, 2, and 4 mM). Coagulation is assessed by the inability of the sample to flow when the tube is inverted. PS-NPs induce complete coagulation at 4 mM Ca²⁺. (D-E) Quantification of fibrin clot properties in the presence of microplastics. (D) Arbitrary size (AU) of fibrin clots. (E) Weight (mg) of fibrin clots. Data are shown as mean ± SEM. *: p < 0.05, ****: p < 0.0001.

The resultant clots were morphologically robust and indistinguishable from positive controls, indicating that PS-NPs support the formation of structurally stable thrombi(Figure 4C-E). These findings extend the observation of platelet activation to a full pro-thrombotic outcome, where PS-NPs act as critical structural components of the thrombus^5, 24^.

### Microscopic Confirmation of Rapid Nanoplastic Incorporation into Platelets

To visually confirm the rapid uptake kinetics observed by flow cytometry, we employed fluorescence microscopy. PS-NPs association with platelets was evident as early as 1 minute post-exposure, with the fluorescence signal becoming increasingly widespread and intense over time (Figure 5). This visual confirmation aligns with flow cytometry data, which demonstrated near-saturation of MFI by 10 minutes, further supporting the conclusion that PS-NPs internalization occurs almost instantaneously upon contact with platelets^5, 24^.

**Figure 5.**
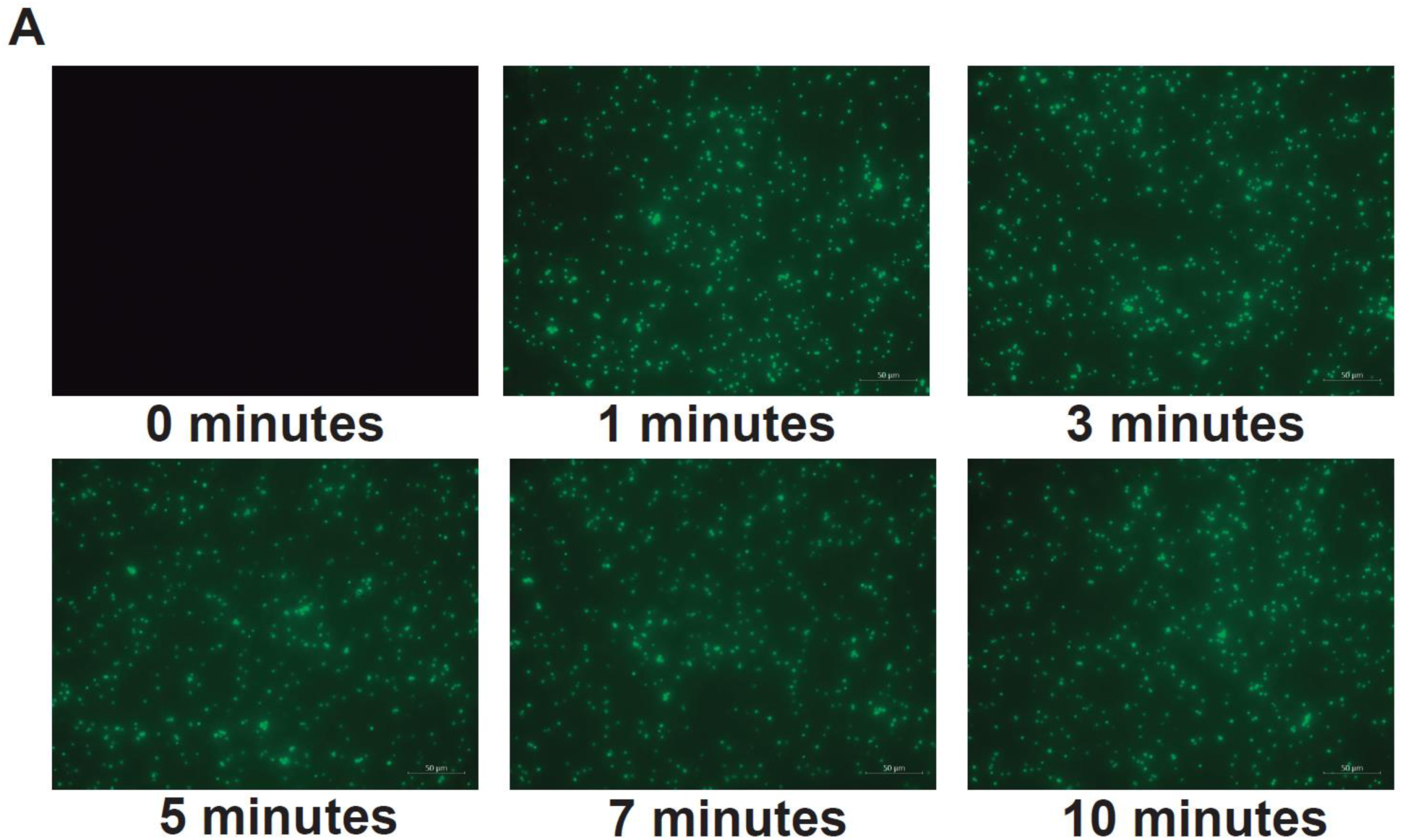
Time-Dependent Polystyrene Nanoplastic (PS-NPs) Incorporation into Washed Platelets. (A) Fluorescence microscopy images showing the kinetics of PS-NPs uptake by washed human platelets over 10 minutes (0, 1, 3, 5, 7, 10 minutes). The 0-minute image is a negative control showing no background fluorescence. Distinct green fluorescent foci appear rapidly, confirming NP incorporation into platelets. Scale bar = 50 μm.

## Conclusions

Our study demonstrates that PS-NPs rapidly associate with platelets, inducing activation, cytoskeletal remodeling, and morphological changes that resemble responses to strong physiological agonists. Platelets exhibit high-capacity, non-saturable uptake of PS-NPs, indicating that PS-NPs accumulation may scale proportionally with environmental exposure, thereby enhancing platelet activation, aggregation potential, and overall thrombotic propensity. In whole blood, PS-NPs further promote coagulation by integrating into fibrin networks and supporting the formation of stable thrombi. These mechanistic insights provide a biological framework linking environmental PS-NPs exposure to heightened thrombotic risk.

Given the detection of MPs/NPs in human blood^1^ and the central role of platelets in thrombotic disease, our findings identify circulating MPs/NPs as a previously unrecognized environmental risk factor for cardiovascular events. Importantly, our in vitro results align with emerging *in vivo* evidence showing that PS-NPs can drive vascular obstruction through interactions with immune cells, leading to impaired blood flow and thrombus formation^9^. Together, these converging lines of evidence suggest that PS-NP-induced platelet activation may synergize with PS-NP-mediated microvascular obstruction to amplify thrombotic events at the organismal level.

Defining exposure thresholds and establishing the clinical relevance of PS-NPs-driven thrombogenic mechanisms are now critical priorities. Further in vivo studies are urgently needed to clarify dose–response relationships, assess chronic exposure effects, and determine the extent to which environmental MPs contribute to human thrombotic disease^5, 24^.

## Acknowledgments

During the preparation of this work, the authors used ChatGPT to check the grammar of the writing. After using this tool/service, the authors reviewed and edited the content as needed and take full responsibility for the content of the publication.

## List of Abbreviations

AA: acrylamide A
BPA: bisphenol A
CVD: cardiovascular diseases
FSC: forward scatter
MFI: mean fluorescence intensity
MPs: microplastics
NPs: nanoplastics
PRP: platelet-rich plasma
PS-NPs: Polystyrene nanoparticles
ROS: reactive oxygen species
SSC: side scatter

